# QuantEM: An optimized platform of vision transformer-based models for segmentation and analysis of electron microscopy data

**DOI:** 10.64898/2026.08.06.743293

**Authors:** Christopher Acree, Evan Krystofiak, Kathryn Coate, Kathleen DelGiorno, Nathan Winn, Sammy Weiser Novak, Elma Zaganjor, Mark Magnuson, Rafael Arrojo e Drigo

## Abstract

Electron microscopy (EM) is essential for resolving cellular ultrastructure, yet quantitative analysis remains limited by labor-intensive segmentation and the scarcity of generalizable models. Here we present QuantEM, an open-source platform for segmentation and analysis of EM data across imaging modalities, tissues, and species. We assembled the largest curated collection of intracellular EM datasets to date, comprising over 15,000 two-dimensional images and 1,700 three-dimensional acquisitions from more than 600 datasets, including nearly 4,000 newly released acquisitions. Using this resource, we trained an EM-specific vision transformer foundation model and systematically optimized adaptation strategies for organelle segmentation. QuantEM provides pretrained models for mitochondria, endoplasmic reticulum, nuclei, and lipid droplets, integrated with interactive proofreading and downstream quantitative analyses through standalone and napari interfaces. Across diverse naive datasets, QuantEM consistently matches or exceeds existing models on zero-shot segmentation while requiring less data for finetuning. We further demonstrate its utility by revealing previously unrecognized subcellular compartmentation of hepatic glucokinase using immuno-electron microscopy.

## Introduction

Electron microscopy (EM) has played a foundational role in the birth of modern cell biology, as it has allowed for the discovery of the constitution of the cell and of different organelles and their function, including the endoplasmic reticulum (ER) and the secretory machinery, mitochondria, and the cytoskeleton^1–4^. Recent technological advancements have pushed working EM resolutions and introduced various forms of volumetric imaging for 3-dimensional (3D) reconstruction and higher sample throughput^5–7^. Together, these changes are generating increasingly larger images and datasets that require next-generation approaches for comprehensive image analysis and reconstruction. Quantitative EM image analysis includes measurements of cellular composition, organelle morphology, localization, and contact sites – all of which are important metrics to characterize changes to cell structure and function in health and disease. However, these analyses typically require segmentation, or the categorization of images into object classes or categories, which is often achieved via manual annotation of EM images. This necessary step involves laborious time commitments from users and can consume days to several months and it is essential to several EM analysis workflows^8,9^. As such, there is a pressing need for the development of intelligent and automated segmentation that allow for a greater range of analyses to match the increased throughput of modern EM.

Over the past decade, there have been increasing attempts to incorporate automation to segment EM images, particularly for mitochondria^10–19^. The recent expansion of deep learning architectures has led to breakthroughs in artificial intelligence (AI) capabilities in microscopy and other types of imaging including the satellite and medical fields^20–22^. However, segmentation of EM images remains challenging due to the diversity of biological and imaging contexts. Many studies continue to use either manual segmentation or to train custom models for their datasets to achieve the required accuracy for their analyses^13,23–25^. Along these lines, we have recently developed custom 2D U-Nets to achieve efficient organelle segmentation, including mitochondria and ER^26^, however these models are specific to our datasets generated in house and are likely to underperform on external datasets. Therefore, we sought to develop a new machinelearning approach capable of general-purpose organelle segmentation and that would outperform previously existing models.

Here, we introduce QuantEM, a software package with segmentation models and an optimized fine-tuning strategy for EM image segmentation and analysis at scale. QuantEM includes models for nuclei, mitochondria, ER, and lipid droplets in two sizes to accommodate variable system requirements, with built-in tools for segmentation proofreading and common analyses. We also are releasing a directory of EM images containing intracellular structures and volumes, including 14,578 assets from public sources and 3,249 newly available assets. Our best-performing models achieve an average Dice of 0.630 for mitochondria, 0.725 for nuclei, 0.453 for ER, and 0.577 for lipid droplets (LDs) across diverse datasets. Moreover, we have determined the best areas and optimization strategies to maximize the performance of pre-existing algorithms to significantly increase segmentation accuracy, while exposing a guided fine-tuning workflow to allow users to adapt our models for custom imaging conditions using minimal annotations. Finally, as a proof of principle, we show how QuantEM can be applied to perform segmentation of challenging EM datasets to map the spatial architecture of cells.

Therefore, our approach introduces an optimized, validated segmentation framework and a rich data resource for development of EM segmentation pipelines and investigation of organelle and subcellular compartmentation patterns at scale.

## Results

### A Unified Directory of Public Intracellular Electron Microscopy Images

The rise of transformer architectures and large language models (LLMs) have shown the ability of modern AI methods and datasets to improve automated analyses. Therefore, we set out to investigate ways to improve upon existing deep-learning (DL) pipelines and to create our own platform to enhance segmentation of large EM datasets. Of note, most deep-learning based EM analyses are trained on similar datasets, largely due to the relatively low abundance of public datasets available as compared to other imaging modalities used to train large vision models. While dedicated repositories and an increasing trend towards open science and reproducibility have started to take root, public EM datasets remain sparse and fragmented. We hypothesized that these limitations significantly compromise the development of effective EM image segmentation algorithms and that increasing available datasets and widening image heterogeneity would be beneficial to the EM field and yield better results.

To address this gap in the field, we took three approaches: first, we set out to create a unified catalog of all known public EM datasets in cell biology; second, we started a global outreach initiative to collect images from previously published works and, third, we compiled a new release of previously unpublished EM images from our group and collaborators (**Fig 1A**). To compile a unified catalog of EM images, we developed code to search all potential data entries in both established image-specific and general data repositories, namely WebKnossos, BossDB, OpenOrganelle, EMPIAR, BioImage Archive, Zenodo, Dryad, and FigShare. Next, we used an LLM-based pipeline to review their metadata, filenames, and associated publications and figures to assess whether they contained intracellular EM data (**Fig 1A-B**). As our goal was to optimize ultrastructural analysis and segmentation of the intracellular space, we excluded images of surface morphology, viral particles, and other image types not containing biological intracellular structures. After extracting the relevant images, we cataloged them based on the available information to identify n=501 unique datasets with n=12,863 unique 2D images and n=1,715 3D acquisitions^30–69^.

**Figure 1.**
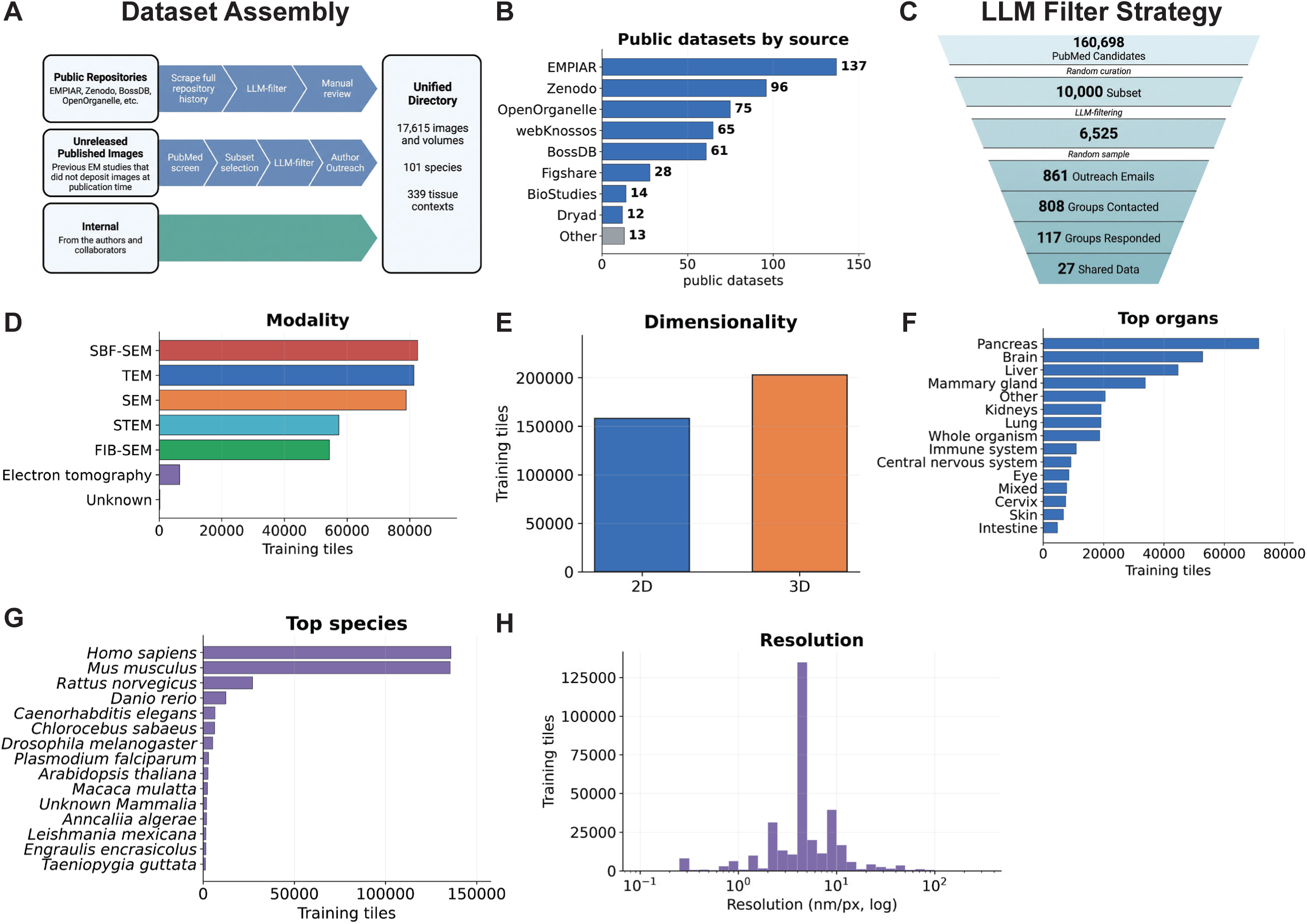
Dataset assembly and data composition. **(A)** Schematic illustration of the strategy used to assemble the electron microscopy (EM) image dataset for this manuscript. **(B)** List of public image repositories used as data source and the number of EM datasets obtained from each respective repository. **(C)** Filtering strategy for EM study identification, author outreach, and addition of new EM data to public repositories. **(D-H)** Composition of the training corpus, tile-weighted, by (D) EM imaging modality, (E) dataset dimensionality, (F) organ, (G) species, and (H) resolution (nm/px) in log scale.

To complement this initial approach, we next ran a PubMed search for any studies published since 2011 containing keywords indicative of EM imaging and identified approximately 160,000 candidate EM studies. Next, we created a subset of 10,000 studies and developed an LLM-based pipeline to determine if the study used EM imaging and did not have the original images available in a repository or linked file. Studies labelled with non-accessible EM data were categorized by corresponding author and research group. We then contacted n=861 of these corresponding authors with personalized outreach communication emails requesting that they provide the EM data from their published works to be deposited to a public repository (EMBL’s BioImage Archive) and to our study. After correcting for bounced emails, we successfully emailed 808 groups, of which 117 responded to our initial outreach and n=27 ultimately contributed with original images from their publications^30–69^ (**Fig 1C, Fig S1A, Supp. Table 1**). In addition to these images, we are releasing a new set of previously unpublished image assets gathered by our group and collaborators. In total, this effort resulted in the release of n=3,249 new images and volumes across n=154 datasets to the research community, an increase in the diversity of the total publicly available EM corpus by ∼30% (**Fig S1B**). The final dataset from all sources encompasses n=101 species, n=339 distinct biological tissues, n=16,107 2D images and n=1,720 3D data acquisitions (**Fig S1C-G**). This complete dataset is publicly available and organized in a browsable directory (See **Data Availability**).

To optimize use of the compiled dataset for training of a new foundation model, we developed an image tiling strategy that created minimally overlapping 2048×2048 tiles from all assets and applied a simple filter to check for a majority of the tile containing information on biological tissue or material (e.g., cytoplasm, nucleus, organelles and avoid blank spaces). During training, these large tiles are randomly kept at native resolution or downscaled at up to 4x, and then randomly cropped to the training model’s context size, after which common augmentations are applied. We balanced the data sources (derived from multiple tissue types) to prevent large 3D volumes from dominating tile share by capping each individual asset to a maximum of 400 tiles while maintaining their native resolution (**Fig 1D-H, S1H-I, Supp. Table 2**). This strategy provides nearly unlimited view diversity from the same base set of images, reducing the risk of model memorization of training data.

### Selection and optimization of a training strategy for EM foundation pretraining

Next, we created broadly applicable organelle segmentation models using a multi-step framework, composed of ***i)*** a new foundational EM image model to encode dense representations of EM images, ***ii)*** decoder heads and adapters which could utilize those representations in diverse segmentation tasks, and ***iii)*** test-time adaptations and fine-tuning (**Fig 2A**). To evaluate the training and performance parameters for this new model, we first investigated the architecture for two recently published EM-native models: OmniEM, a vision transformer with large architecture (ViT-L, with 302 million parameters) model based on the DinoV2 architecture, and EMCellFound (EMCF), which used a masked autoencoder approach^70,71^. We tested which self-supervised training strategy produced the most transferable representations for EM images as well as baseline DinoV2 and DinoV3 encoders^72,73^. We excluded Segment Anything (SAM) and SAM-based encoders (e.g. MicroSAM) in this experiment; while these models have proven useful in applications to compact objects, these studies themselves note these models’ deficiencies in fine structures and thin networks^16,74,75^. Therefore, since ER segmentation was a primary target of our study and the need for accurate segmentation of other network-like structures (e.g., plasma membrane), we prioritized models more suited to develop and apply these capabilities. Since segmentation requires pixel-accurate delineation of membranes, organelles, and thin cellular protrusions, we did not evaluate models using pretraining loss alone. Instead, each candidate encoder was evaluated using a fixed frozen-backbone protocol in which identical light-weight UPerNet decoders were trained on ER and mitochondria segmentation tasks and benchmarked against ground truth ER and mitochondria masks that were generated via manual annotation (**Fig 2B**). Here, OmniEM only modestly outperformed its generalist equivalent DinoV2, and the more recent model DinoV3 matched or exceeded it in both tasks when frozen (**Fig 2B-C**). Given this initial result, we ran the same test using a Low-Rank Adaptation (LoRA) approach, which significantly increased OmniEM performance versus the base Dino models and EMCF^76^ (**Fig 2B-C**).

**Figure 2.**
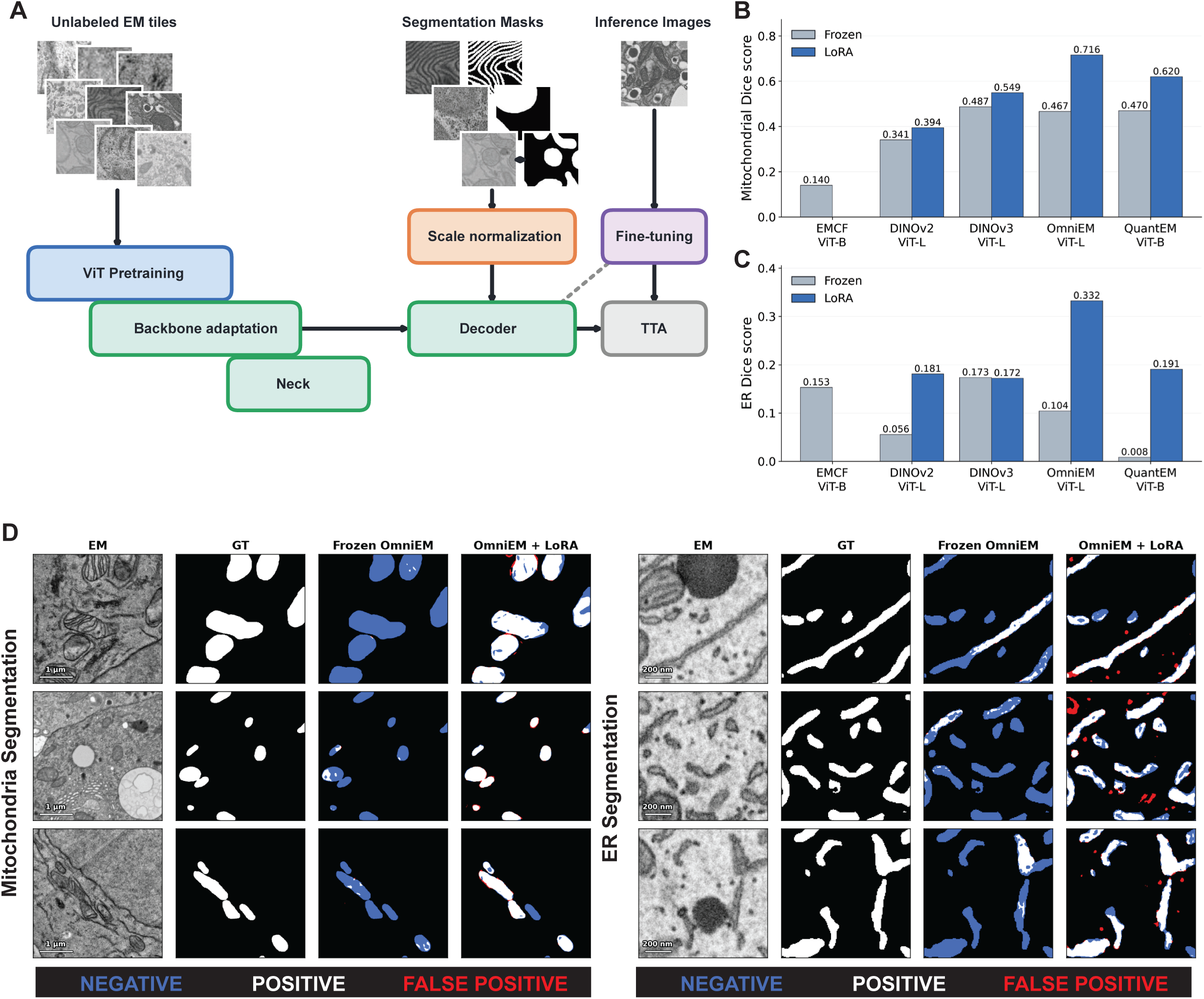
Foundation model pretraining. **(A)** Schematic showing the different architectural components trained and tested for our organelle segmentation models. **(B)** Mitochondria segmentation Dice scores for generalist and EM-native models evaluated on the same test dataset, under both frozen backbone and LoRA adaptation conditions. **(C)** Same as B for ER segmentation Dice scores. **(D)** Representative images showing EM images, ground-truth manual segmentations, the frozen OmniEM backbone’s segmentation performance using a UPerNet on mitochondria and ER segmentation data, and the LoRA adapted OmniEM’s segmentation performance under the same decoder and dataset.

Based on the increase in base performance between base DinoV2-3, and between base DinoV2 and OmniEM after LoRA, we tested the robustness of our dataset and tiling approach by training a new ViT-B model (86 million parameters) based on the DinoV3 architecture (called QuantEM). Pilot experiments revealed that Gram anchoring and context windows above 512px did not improve segmentation capabilities during early training, so we excluded them to increase training speed (**Fig S2A-B**). After an initial training ramp, we hypothesized that the incorporation of metadata into the encoder might additionally boost training and performance results. We tested whether the incorporation of scale and modality metadata into pretraining using a “FIne tuning with NO labels” (FINO) method, which has been recently proposed to adapt models to specialized domains through the boosting or suppression of defined metadata factors^77^. This approach, however, did not yield improvements over the model baseline and was marked by significant discrepancies between ER and mitochondria tasks (**Fig S2C-D**). As such, we continued training under our original training regime.

Next, we focused on the performance of our newly trained model QuantEM, which showed continued improvements through ∼650,000 training steps before plateauing on segmentation performance (**Fig S2E**). We observed similarly large responses to backbone adaptation as OmniEM, and its final Dice scores in the fixed decoder test on the adapted backbone were comparable with OmniEM for mitochondria, outperforming all other non-domain native equivalent models for both organelles (**Fig S2F**). Importantly, this performance means that QuantEM with adaptation can achieve increased segmentation accuracy of mitochondria and ER compartments in naïve datasets (**Fig 2D, Fig S2G**). This also indicates that our ViT-B model attained similar performance to the significantly larger ViT-L Omni-EM despite having approximately a third of the parameters and associated training workload. From these results, we continued our experiments using OmniEM as a ViT-L base model and QuantEM as our ViT-B base model.

### Creating Decoders for EM Segmentation Tasks

Next, we sought to determine what architectures would result in the best segmentation performance for mitochondria and ER compartments. To do this, we experimented with different options of necks, decoders, loss functions, levels of backbone adaptation, input standardization, image style-conditioning, multi-organelle vs single-organelle decoders, and test-time adaptation. Here, we isolated and evaluated each of these parameters separately, finding that backbone adaptation, input scaling, loss function, and decoder choices all had large impacts on segmentation performance, while test-time adaptation, dataset modulation, image-style conditioning, neck injection, and multiscale models had more modest to negligible effects (**Fig 3A-B, Fig S3A-G and -F)**. Here, backbone adaptation mechanisms were by far the most impactful lever on segmentation performance; whereas LoRA was the best adaptation mode for most ViT-L models, ViT-B models benefited more from deeper adaptations of the last 4 transformer blocks and full fine-tuning (**Fig S3A-B**). This was despite the lower learning rates for these adaptations as used in our setup, as we wanted to test adaptation on a fixed step budget, and higher learning rates during fine-tuning caused model instability and collapse^78^ (**Fig S3C**).

**Figure 3.**
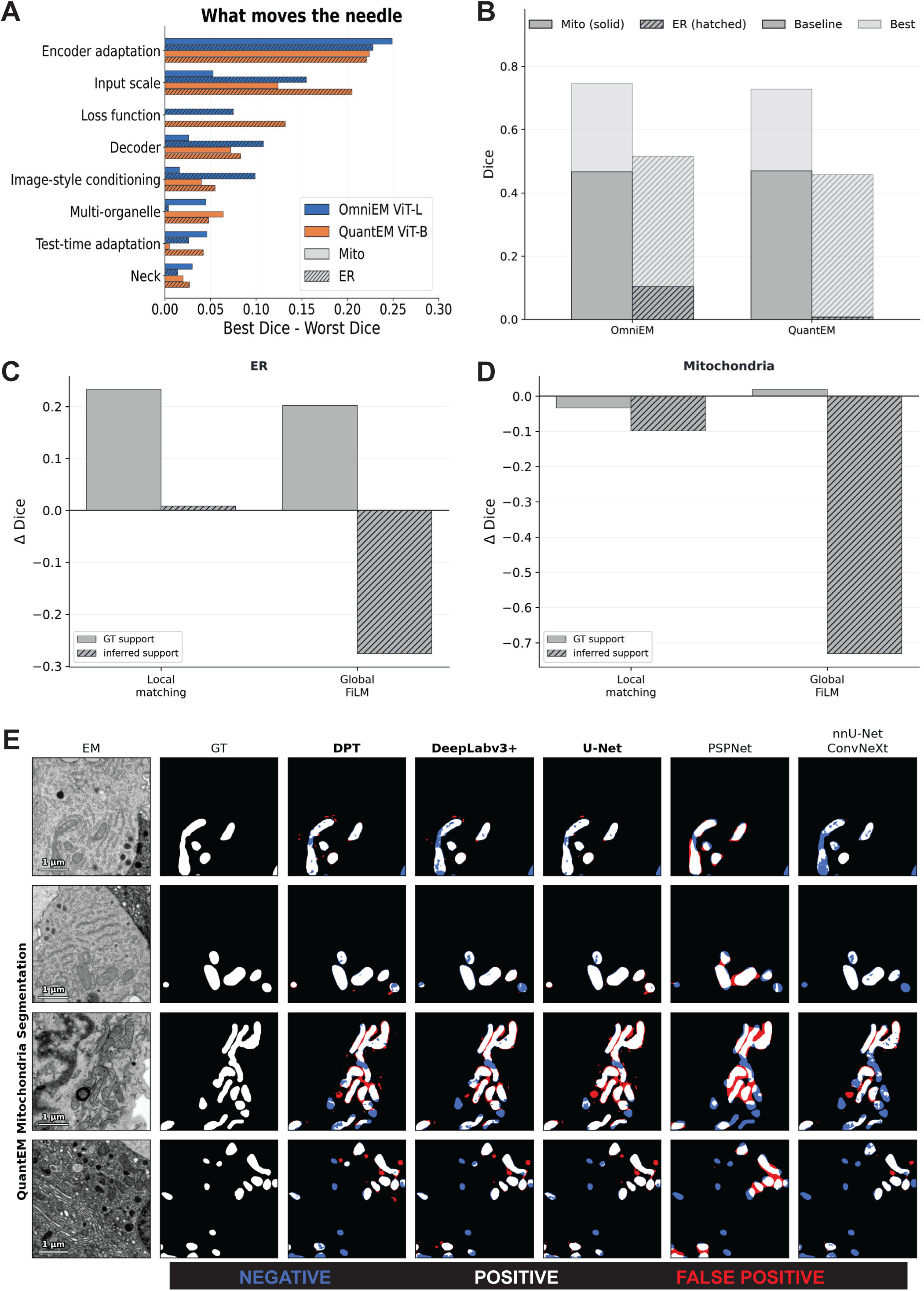
Optimizing EM organelle segmentation. **(A)** A list of parameters for which we ran experiments testing segmentation performance under otherwise identical configurations, and the associated mitochondrial and ER Dice coefficient gaps between the best and worst performing variants in that experiment. **(B)** Dice coefficients for ER and mitochondria segmentation tasks under baseline frozen backbone conditions for each encoder model and using the best configuration combination tested. **(C)** Change in Dice coefficient for ER segmentation when a LoRA-adapted OmniEM ViT-L is given test-time support, relative to the same model with support disabled. Pixel matching builds foreground**/**background prototypes from the support region and re-scores every pixel by cosine similarity^102^; Pooled global instead average-pools the support features into a single appearance code that conditions the decoder by Feature-wise Linear Modulation (FiLM) before re-segmenting. GT = support taken from manual annotations; inferred = support taken from the model’s own first-pass predictions. **(D)** Differences in mitochondria segmentation, as with (C). **(E)** Representative images of outputs from mitochondrial segmentation decoders trained with the given architectures on the same QuantEM encoder backbone.

Next, we hypothesized that having a fixed higher resolution for ER would lead to improved performance due to its thin, curvilinear structure. Our initial baselines accordingly used a fixed 2nm scale; however, we found fixed 4nm and 8nm resolutions both outperformed the 2nm scale and neither had a significantly different Dice than a decoder trained and run at native resolutions (**Fig S3D**). For mitochondria, the fixed 8nm and native resolution trained decoders outperformed 4nm and 16nm fixed decoders on Dice (**Fig S3E)**. Similarly, we observed minimal mitochondria segmentation changes from averaging multiscale inputs (0.75x, 1x, 1.5x) or from providing dual-scaled inputs to the attention mechanism^79^, while these modifications actively harmed ER segmentation (**Fig S3F-G**). We next tested if “positive” examples could boost segmentation performance at test-time. We evaluated both a pixel similarity approach and global FiLM conditioning based on provided foreground/background examples from either ground-truth (GT) annotations or from select first-pass predictions. While mitochondria showed only slight differences from its stronger base model, ER segmentation was significantly enhanced when GT examples were provided^78,80^. However, attempting to infer organelle support examples from automatically selected organelle segmentation predictions could not recapitulate high-accuracy gains in segmentation (**Fig 3C-D**). Of note, decoder, neck, loss function modification, and image-style encoding experiments showed relatively small differences between top configurations, with significantly inferior performance only for the worst options (**Fig S4A-F**)^78,81–92^.

### Final Model Performance and real-world application of QuantEM

To assess the generalization performance of QuantEM, we assembled manually annotated public datasets for mitochondria, ER, nuclei, and lipid droplets together with annotated in-house images, and created orga-nelle-specific training sets with “held-out” test sources. We trained the nuclei and lipid droplet models using the mitochondria-derived architecture, while standardizing the input resolution to 8nm for lipid droplets and 25nm for nuclei. We compared optimized QuantEM and OmniEM models against publicly available organelle and EM segmentation models. Across diverse unseen datasets, QuantEM and OmniEM consistently matched or exceeded existing methods, achieving the largest improvements for ER segmentation while matching best-in-class model performance for mitochondria and lipid droplets (**Fig S5A-D**). To directly compare against specialized models, we reconstructed the published train/test splits for OrgSegNet and DeepContact within our benchmarking framework. OmniEM and QuantEM models were competitive with OrgSegNet on its native plant nucleus and mitochondria dataset and substantially outperformed DeepContact on its native COS-7 cultured cell dataset while maintaining superior performance across unrelated biological datasets, demonstrating robust generalization beyond the original training domains (**Fig S5A-C, S6A-B**).

To evaluate QuantEM under particularly challenging imaging conditions, we next applied it to immuno-electron microscopy (ImmunoEM). Unlike conventional EM, ImmunoEM sample preparation must simultaneously preserve ultrastructure and antigen accessibility, often reducing membrane contrast, making automated organelle segmentation substantially more difficult and thus making immuno-EM one of the most challenging settings for automated EM segmentation. We therefore used ImmunoEM as a stringent benchmark for model adaptation and down-stream quantitative analysis. We analyzed liver sections from mice subjected to fasting and glucose infusion and manually annotated a small number of representative regions to evaluate adaptation efficiency. As expected, baseline performance was insufficient for all models, especially MitoNet, because ImmunoEM images exhibit reduced membrane contrast relative to conventional EM. Remarkably, fine-tuning with only two manually annotated image crops increased QuantEM and OmniEM performance to approximately 87% Dice, whereas MitoNet required substantially more annotations, never achieved the same peak accuracy, and exhibited overfitting as additional labels were added (**Fig S6C**). Notably, QuantEM and OmniEM achieved this performance by updating only lightweight decoder heads, reducing both the number of trainable parameters and computational cost relative to full-model fine-tuning.

Having established accurate segmentation performance, we next used QuantEM to quantify the intracellular distribution of glucokinase (GCK) in hepatocytes. GCK translocates from the nucleus to the cytoplasm following glucose stimulation, but the spatial organization of cytoplasmic GCK remains poorly understood^93–96^. Combining automated organelle segmentation with correlative MIMS-EM enabled automatic assignment of gold-labeled GCK molecules to intracellular compartments. QuantEM accurately recapitulated the expected nuclear localization of GCK during fasting while enabling quantitative spatial analysis of its cytoplasmic distribution following glucose stimulation **(Fig. 4C)**. Cytoplasmic GCK was significantly enriched both within mitochondria and within a 100-nm mitochondrial neighborhood compared with randomized particle distributions generated by Monte Carlo simulation in the 6hr fast and glucose-infused conditions **(Fig. 4D)**. The higher levels of nuclear gold enrichment combined with higher percentage of cytosol occupied by mitochondria in the overnight fasting condition may explain the lack of observed mitochondrial enrichment of cytosolic GCK compared to random. Nevertheless, the majority of cytoplasmic GCK remained outside these regions, indicating that mitochondrial association represents one component of a broader cytoplasmic distribution.

**Figure 4.**
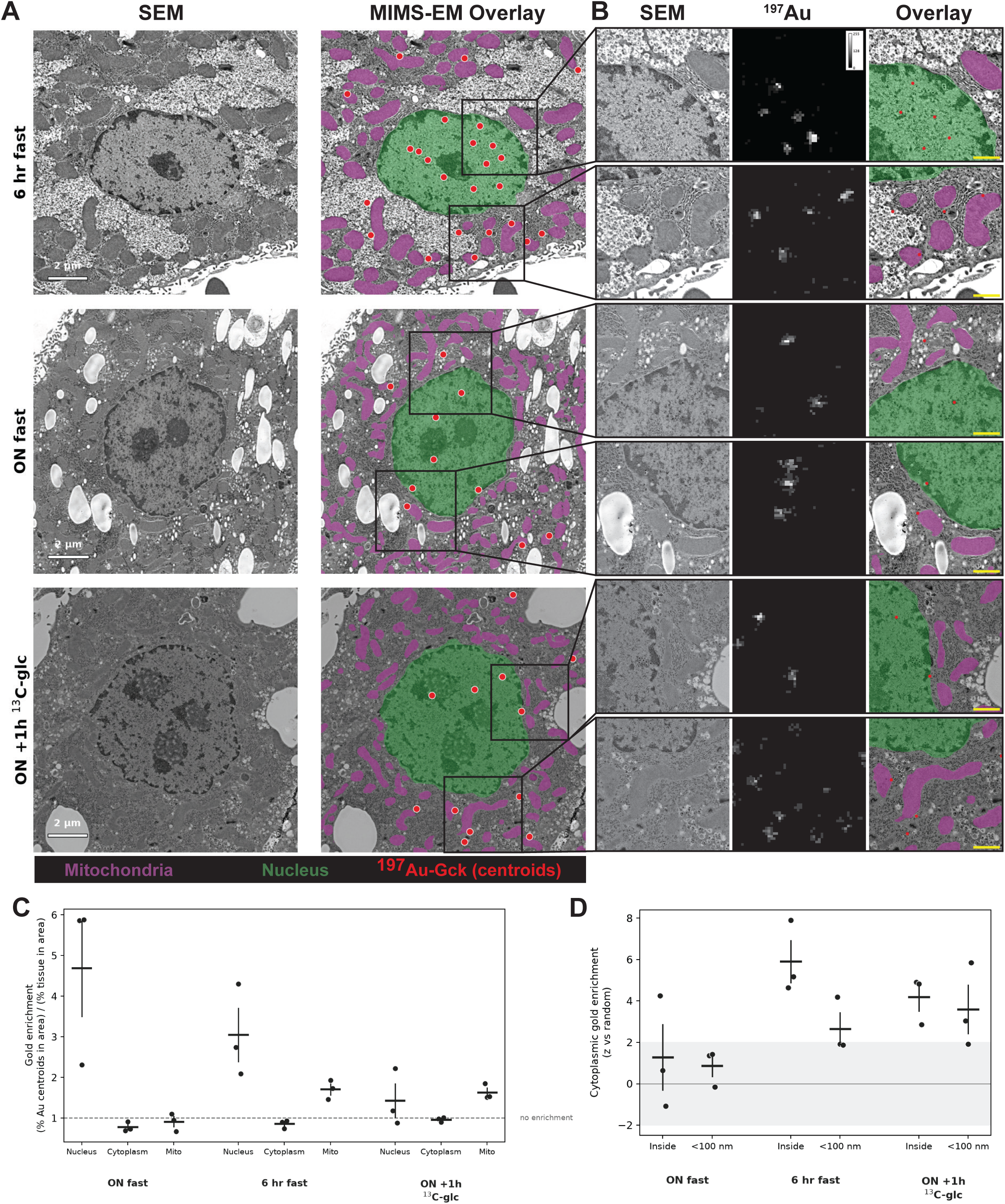
ImmunoEM segmentation and Gck localization. **(A)** Representative EM images in each experimental group. Left, raw EM image; right, EM overlayed with QuantEM organelle segmentation and ^197^Au centroids. **(B)** Two zoomed-in sections of the same crops from (A), with raw ^197^Au signal included. **(C)** Gold point enrichment in each compartment with gold-counts normalized to the percentage of tissue containing that compartment. ON fast=Overnight fast (13 hours). ON +1h^13^C-glc=Overnight fast (13 hours) followed by 1 hour of ^13^C-glucose infusion at 40mg/kg-min. **(D)** Cytoplasmic gold enrichment inside mitochondria, or outside of mitochondria but within 100nm of a mitochondrial border, expressed as a z-score against 20 Monte Carlo simulations placing the same number of particles at random within the same tissue mask (z = (observed − null mean) / null s.d.; 0 = chance, grey band |z| < 2). Nuclear gold is excluded from both observed and simulated values, so this is a within-cytoplasm measure independent of any nuclear enrichment.

Together, these results demonstrate that QuantEM can be rapidly adapted to challenging EM imaging datasets and enables quantitative spatial analyses that would otherwise require extensive manual annotation.

## Discussion

Recent studies have debated the utility of foundation vision models for EM image analysis and whether they offer meaningful advantages over established convolutional architectures such as U-Nets^97,98^. Our results demonstrate that EM-native foundation models can match or exceed the performance of specialized segmentation networks while requiring substantially less task-specific annotation for adaptation. We demonstrate that lightweight adaptation strategies allow for quick fine-tuning to naïve datasets, allowing both optimized QuantEM and OmniEM frameworks to achieve high segmentation accuracy from only a small number of manually annotated examples. These properties make foundation models particularly well suited for heterogeneous imaging conditions usually observed in modern EM workflows.

Here, we provide pretrained segmentation models in both base (ViT-B, ∼86 million parameters) and large (ViT-L, ∼302 million parameters) configurations for mitochondria, ER, nuclei, and LDs, together with interactive proofreading and quantitative analysis tools available as both a standalone application and a napari plugin. To benchmark our software, we performed systematic evaluations of the types of architectural and training decisions underlying EM foundation models, and identified backbone adaptation (e.g., LoRA) as the dominant determinant of down-stream segmentation performance. We anticipate that these observations will help guide development strategies for future EM-specific segmentation models.

Although QuantEM consistently matched or exceeded existing methods across diverse datasets, segmentation performance remains substantially below what can be achieved via expert manual segmentation and versus many other natural-image applications^99,100^. This limitation is particularly evident for ER, where thin and morphologically heterogeneous structures present one of the most difficult segmentation targets in EM datasets. While previous work has suggested that limited receptive field and global context contribute to this challenge^100,101^, our results suggest that an enlarged field of view does not help ER segmentation **(Fig. S3F)** and that biological heterogeneity and the scarcity of diverse annotated training data are likely to be more important constraints. Therefore, continued expansion of publicly available EM datasets and annotations are likely to yield further improvements in automated segmentation.

Finally, this study focused exclusively on two-dimensional EM segmentation, reflecting the current need for robust and broadly applicable tools for conventional EM datasets. We expect many of the training strategies and adaptation approaches developed here to translate naturally to volumetric imaging, although optimal architectures for three-dimensional segmentation remain an active area of research. QuantEM provides an extensible framework in which these future advances can be readily incorporated while offering an immediately usable platform for quantitative ultrastructural analysis.

### Limitations of this study

Although QuantEM substantially improves the accuracy and training framework for EM image segmentation and analysis, several limitations remain. First, our models focus exclusively on two-dimensional EM images. While many of the optimization strategies described here are expected to translate to volumetric datasets, segmentation of large three-dimensional EM volumes presents distinct computational and architectural challenges that were beyond the scope of this study. We also note that some pretraining strategies tested here may exhibit performance improvements only when added after initial performance has plateaued, as contrasted to their relatively early implementation here. Second, despite compiling a large and curated EM dataset, publicly available annotations remain limited for many organelles, tissues, and imaging modalities. Consequently, segmentation performance varies across structures, with thin and morphologically heterogeneous organelles such as the ER remaining particularly challenging. Third, QuantEM currently provides pretrained models for four types of organelles. While our results in nucleus and lipid droplets demonstrate the capacity for adopting mitochondrial training strategies to other “blobby” organelles, the performance of the QuantEM ViT-B model on LDs indicates further optimization strategies may be needed for new organelles, and this is likely to be more important for organelles with unique image features like Golgi. Finally, while our benchmarking demonstrates robust generalization across diverse datasets, performance on highly specialized sample preparation protocols or imaging modalities may still benefit from lightweight user adaptation, which QuantEM is designed to facilitate to accelerate discovery.

## Author contributions

C.A. conducted all machine learning experiments, data analyses, created figures and wrote the manuscript. E.K., K.C., K.E.D., N.W., S.W.N., E.Z., M.M., and R.AeD., collected data, provided reagents, performed image analysis, and provided valuable manuscript and data analysis feedback. C.A. and R.AeD. conceptualized the project and wrote the manuscript.

## Acknowledgments

This research was supported by recruitment funds from the Vanderbilt’s Department of Molecular Physiology and Biophysics and NIH-NIDDK grant R01DK138141 to R.AeD. C. A. was supported by the Vanderbilt Molecular Endocrinology Training Program grant 5T32 DK07563. Electron microscopy was performed in part by the Vanderbilt Cell Imaging Shared Resource (supported by NIH grants CA68485, DK20593, DK58404, DK59637, EY08126, R24OD037694, S10MH137068, and S10OD028704). The DelGiorno Laboratory was supported by NIH/NIGMS R35GM142709, NIH/NCI P30CA068485, NIH/NCI P50CA236733, NIH/NIDDK P30DK058404, NIH/NIDDK P30DK020593, the American Gastroenterological Association Research Scholar Award (AGA2021-13), The American Cancer Society Research Scholar Grant (ACS1433470). EZ was supported by NIGMS R35GM154684. R.AeD. is the guarantor of this work with full access to all the data in the study and take responsibility for the integrity of the data and the accuracy of the data analysis. All the authors declare no conflicts of interest.

---

**Figure S1. (A)** Group progression rates through the outreach pipeline. **(B)** Counts of existing and newly available EM datasets. **(C-G)** Per-image composition of the corpus, irrespective of size. Images by (C) kingdom, (D) imaging modality, (E) organ, (F) species, and (G) resolution and imaging modality. **(H)** Distribution by modality of unique images and volumes and of final distribution of training tiles after image tiling and capping at 400 tiles per asset. **(I)** Breakdown by dimensionality for the counts of unique images, total pixels, and final number of training tiles after image tiling and capping at 400 tiles per asset.

**Figure S2. (A)** Dice scores for mitochondria segmentation when trained with identical UPerNet decoders and datasets on frozen-backbone encoders. Two seeds were trained to 50,000 steps (batch size 128 = 6.4 million images) with a model context size of 512px. The “With Gram anchoring” configuration was identical to the base configuration except with Gram loss enabled from step 0. The Large Context Full arm was trained at 1024px context size, the Large Context Ramp Light was trained partially at first 512px and then 768px, and Large Context Ramp Heavy was trained sequentially at 512px, then 768px, then 1024px, see code for full configuration details. **(B)** Dice scores for ER segmentation, as with (A). **(C)** Mitochondrial segmentation Dice coefficient after control or FINO pretraining. After an initial 50k steps, pretraining was continued from the same checkpoint for 30k steps under baseline or while preserving or suppressing the indicated metadata factor using FINO. **(D)** ER segmentation Dice coefficient after control or FINO pretraining, as with (C). **(E)** Dice performance for mitochondria and ER segmentation heads trained with LoRA, evaluated every 50k steps. **(F)** Gains in Dice coefficient between decoders trained directly on the final output layer versus those using the optimized backbone adaptation (LoRA or fine-tuning) on the same encoder. **(G)** Label efficiency curves for different encoders evaluated using mitochondrial Dice coefficient, evaluated using subsets of the same mitochondrial training dataset on an identical test dataset.

**Figure S3. (A)** Dice coefficients for otherwise identically trained mitochondria decoders with different levels of backbone adaptation for each encoder. LoRA=Low-Ranked Adaptation, Last-4=the last 4 transformer blocks of the encoder are unfrozen and fine-tuned. Full FT=All encoder parameters are fine-tuned. Adaptation arms were deliberately run at different encoder learning rates (LoRA 1e-3, Last-4 1e-4, Full FT 2e-5) because full fine-tuning is unstable at higher rates, so each arm was trained near its own usable operating point; the training budget was matched at 10k steps across arms. This compares practical implementation tradeoffs rather than matching equivalent learning. **(B)** Dice coefficients for ER decoders with differing backbone adaptation as in (A). **(C)** Dice coefficients for otherwise identically trained mitochondrial decoders with full fine-tuning for the encoder under different learning rates after 15k steps. Higher learning rates exhibited instability and collapse. **(D)** Dice and boundary F1 scores for ER decoders when evaluated on either native resolution or standardized (up sampled / down sampled) resolution training and test data. **(E)** Dice and boundary F1 scores for mitochondria decoders, as in (D). **(F)** Dice coefficients for ER segmentation under three scale strategies. *Native*: model trained and evaluated at native resolution (0.513). *Multiscale*: a 4nm-resolution-trained model evaluated by averaging its predictions across inputs rescaled to 0.75×, 1× and 1.5× (test-time fusion, no retraining; 0.382). *Dual scale*: a model trained on two streams through the shared encoder, one at 4nm resolution and a 4x larger field of view at 8nm resolution fed together via cross-attention (0.425) single fixed seed. **(G)** As in (F), for mitochondria. The multiscale mitochondria model was trained at native resolution, dual-scale was trained via cross-attention at native scale and a co-centered window with 2x coarser pixels to achieve the same 4x larger field of view. *Native* 0.715, *Multiscale* 0.727, *Dual scale* 0.681.

**Figure S4.** Architectural and training-recipe levers are near-null compared with encoder adaptation. All panels report held-out-source test Dice for OmniEM ViT-L and QuantEM ViT-B, each on its own LoRA (OmniEM) or fine-tune (QuantEM) adapted base. Mitochondrial Dice is semantic-foreground quality computed by fixed-threshold connected components, not instance segmentation. **(A)** ER Dice across six dense decoder heads. **(B)** Mitochondria Dice across nine decoder heads. **(C)** Neck ablation: a resnet34 detail-neck versus no neck (naive 1×1 projection) across the model x organelle x decoder matrix. **(D)** ER loss functions, adding topology terms cumulatively to Dice-BCE. **(E, F)** Mitochondrial (E) and ER (F) Dice under three image-style conditioning strategies. *Baseline*: no conditioning. *Inferred style*: a per-tile style code, computed from low-level appearance statistics (intensity percentiles, mean, s.d., local contrast, gradient energy, noise estimate and radial spectral bins), applied by feature-wise linear modulation (FiLM). *MixStyle*: during training, each feature map’s channel-wise mean and s.d. are replaced by a randomly weighted interpolation between its own and those of another image in the same batch. *DSU* (Domain Shifts with Uncertainty): during training, each feature map’s channel-wise mean and s.d. are resampled from Gaussians whose spread is estimated across the batch. MixStyle and DSU were applied with probability 0.5 during training only.

**Figure S5. (A-D)** Organelle segmentation models benchmarked for (A) mitochondria, (B) ER, (C) nuclei, and (D) LDs.

**Figure S6. (A)** Representative images of mitochondria segmentation by optimized OmniEM, QuantEM, and publicly available models. **(B)** Representative images of ER segmentation by optimized OmniEM, QuantEM, and publicly available models. **(C)** Efficiency of labeling for fine-tuning OmniEM, QuantEM, and MitoNet. LoRA showed minimal gains over head-only fine-tuning by OmniEM, while MitoNet’s performance drops at increased labels is indicative of overfitting.

**Supplementary Table 1.** Previously published studies with newly released EM images and/or volumes to public repositories following our outreach campaign.

**Supplementary Table 2.** All datasets in our corpus, with associated image and volume counts, tile counts in our derived training dataset, and public repository URLs.

**Supplementary Table 3**. List of sources and their respective share of ground-truth segmentation data used in training and benchmarks for each organelle.

