## Supplementary figures and images for "QuantEM: An optimized platform of vision transformer-based models for segmentation and analysis of electron microscopy data"

### Supplementary Figure 1

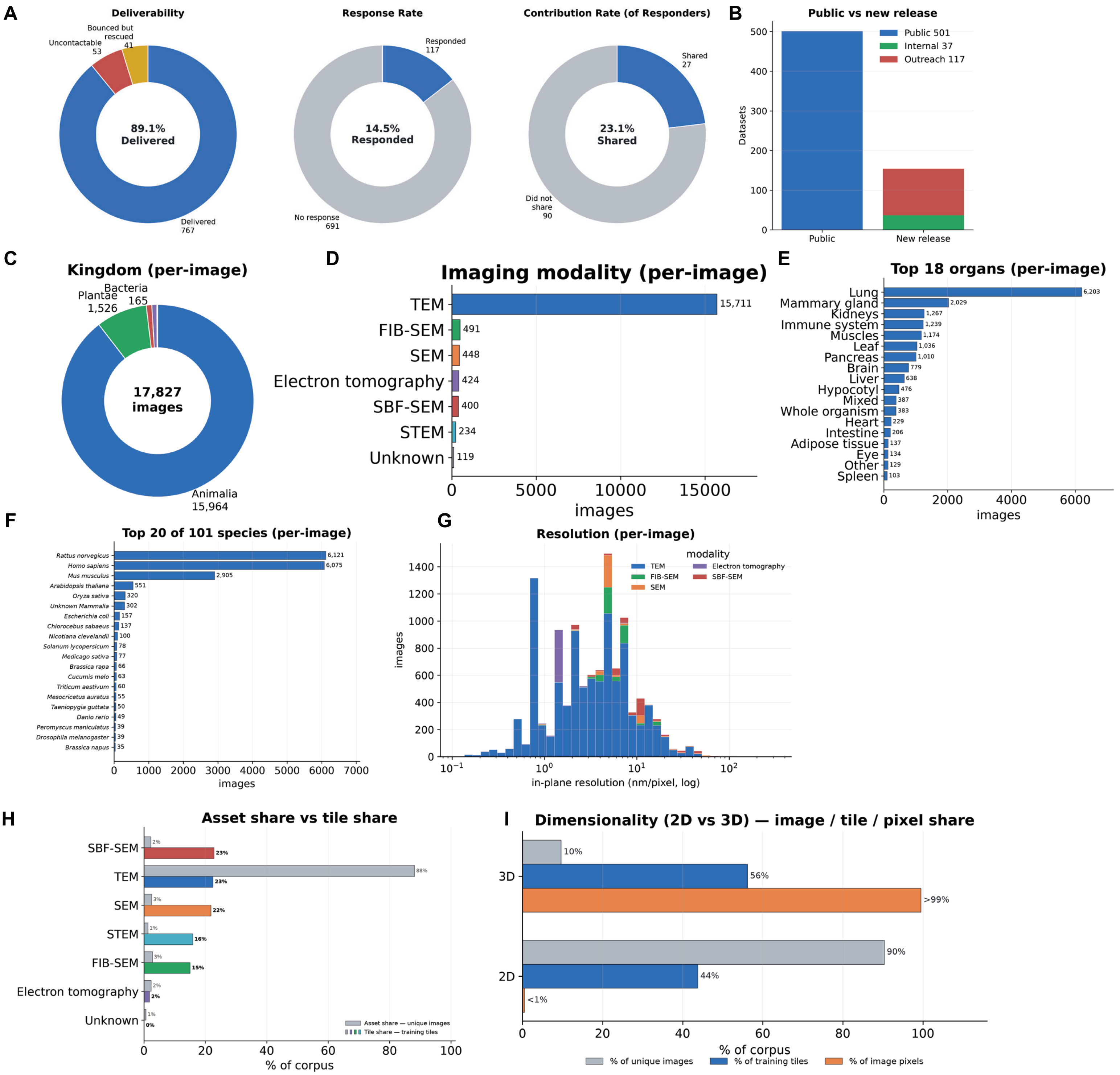

### Supplementary Figure 2

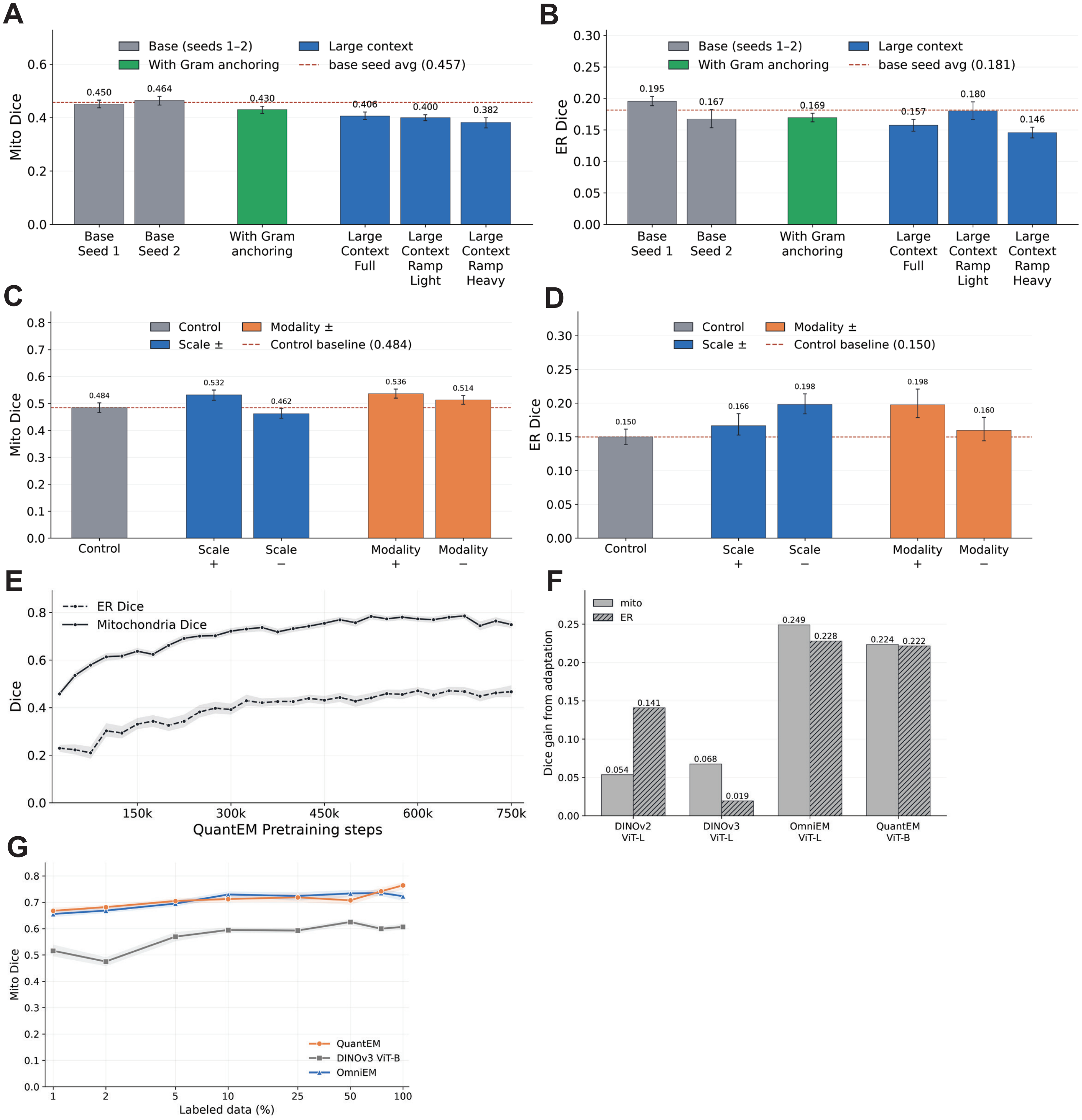

### Supplementary Figure 3

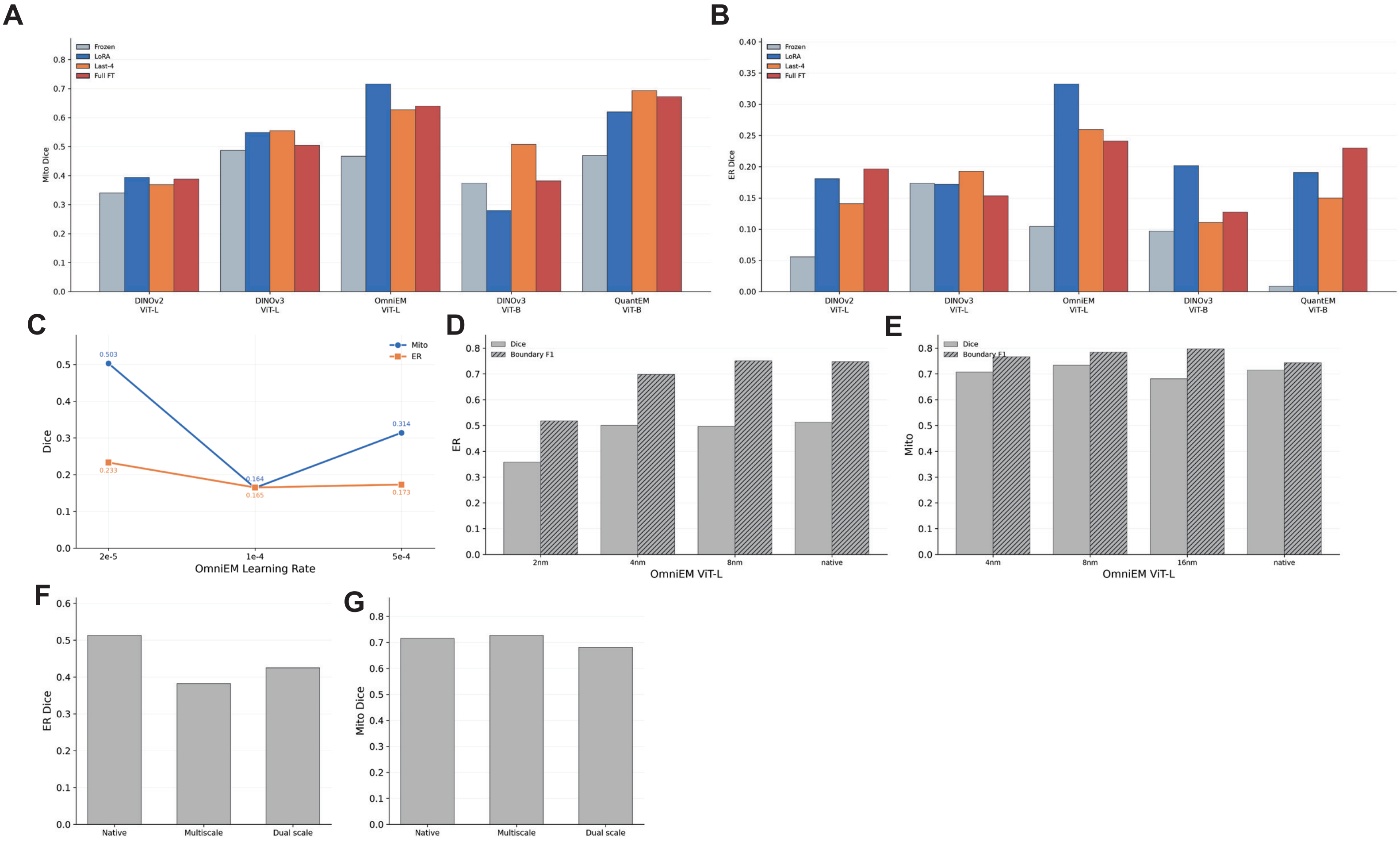

### Supplementary Figure 4

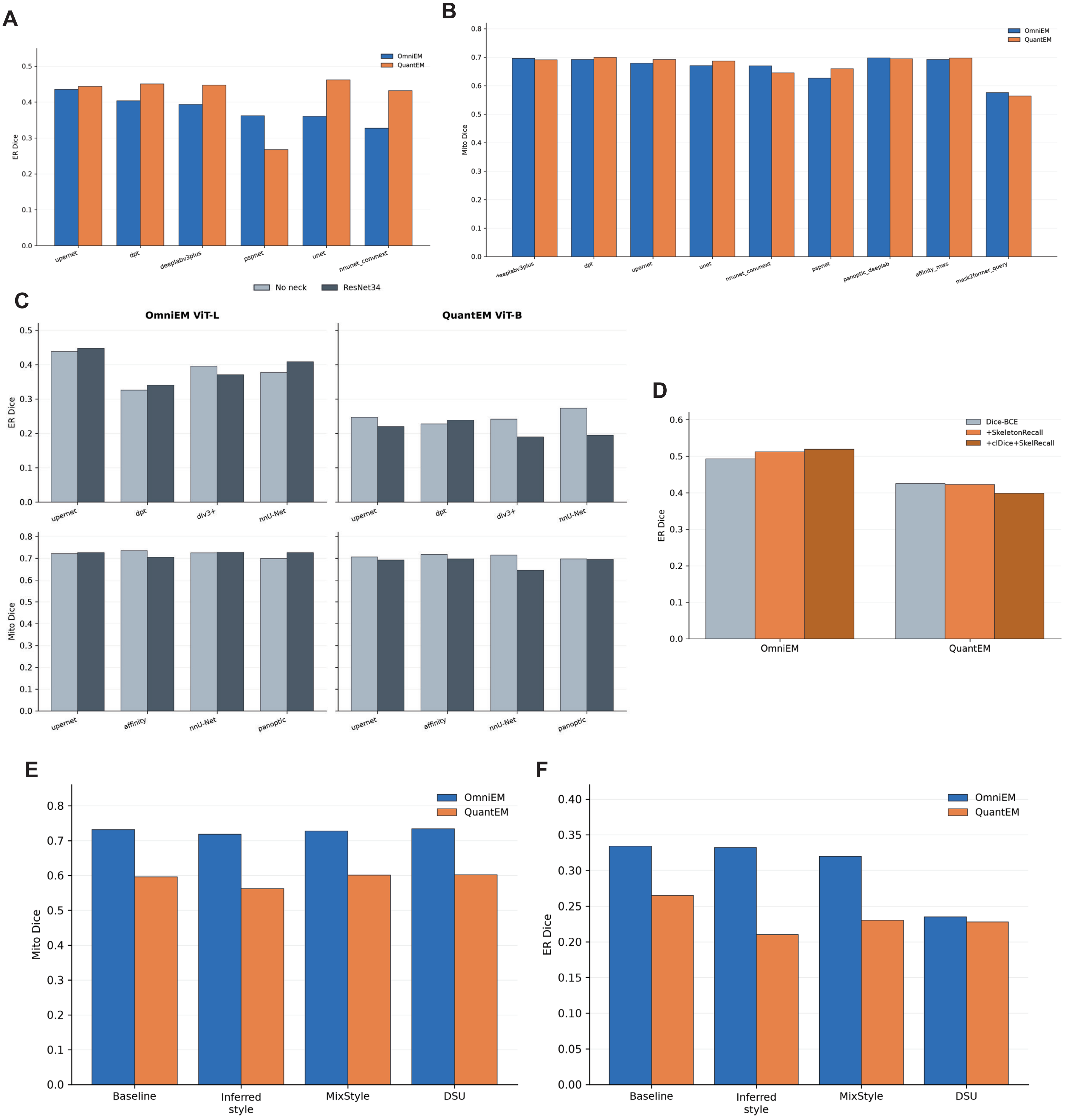

### Supplementary Figure 5

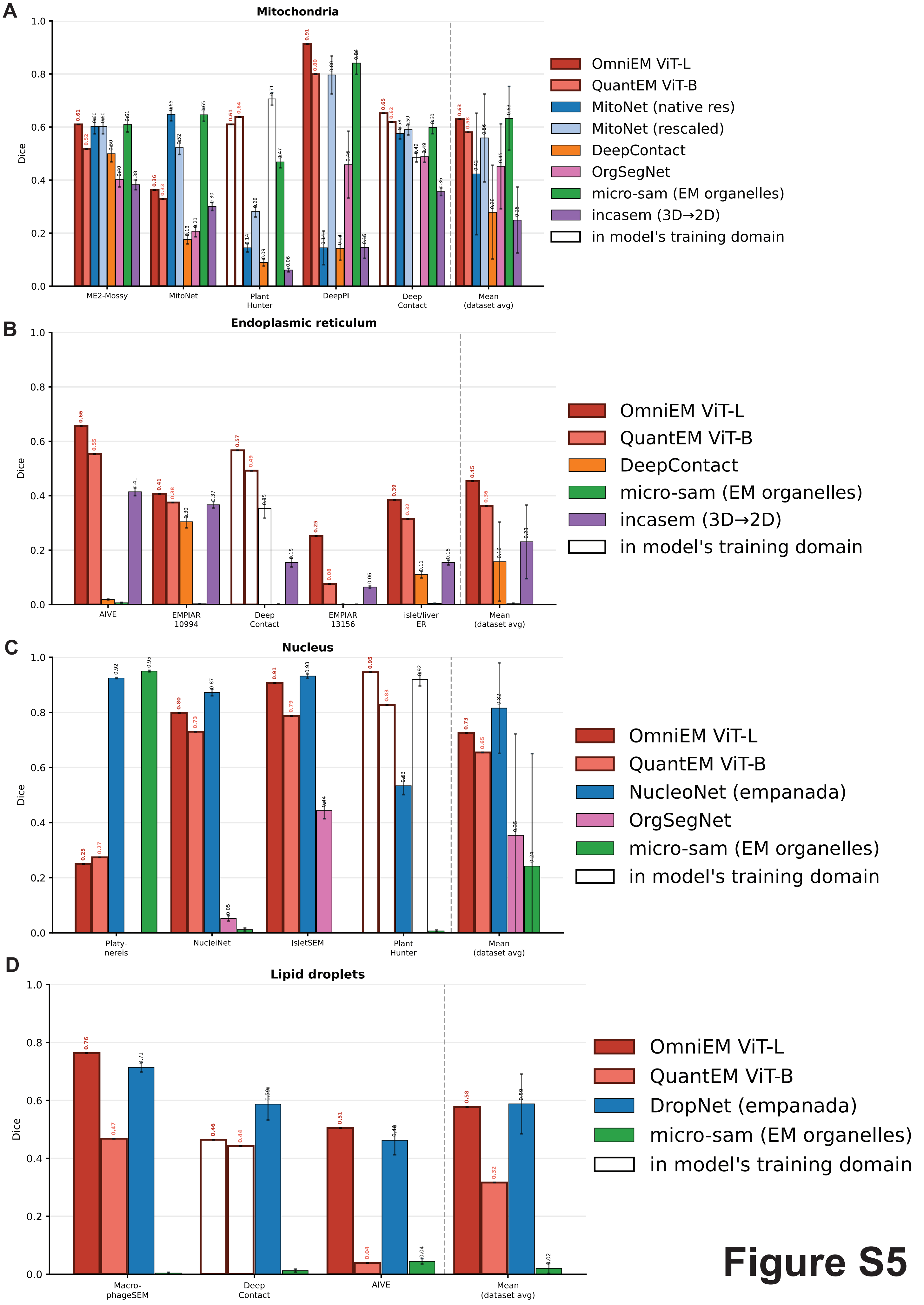

### Supplementary Figure 6

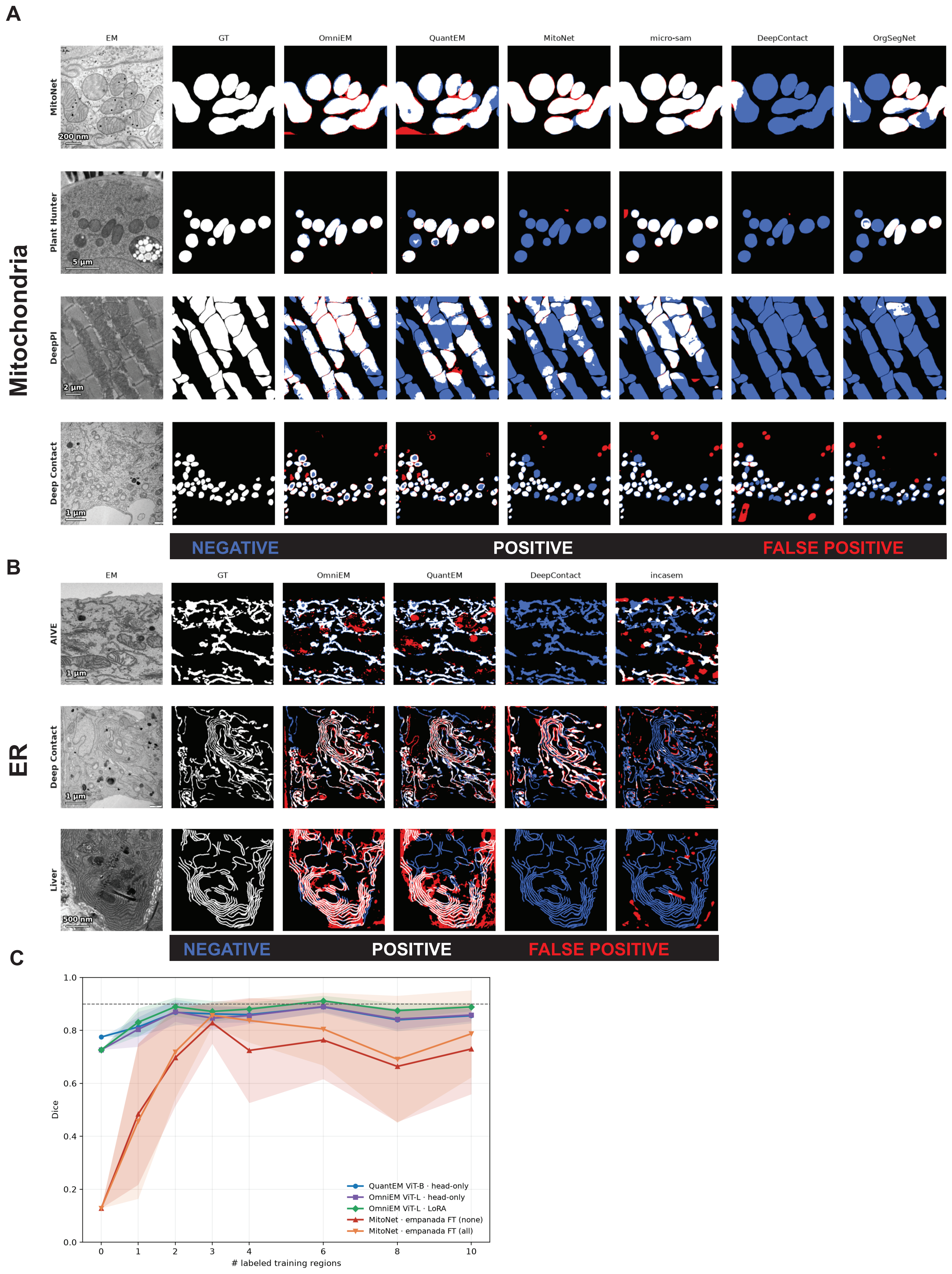
